# Linking Brain Entropy to Molecular and Cellular Architecture in Psychosis

**DOI:** 10.1101/2025.05.08.652921

**Authors:** Qiang Li, Jingyu Liu, Vince D. Calhoun

## Abstract

Brain entropy reflects the complexity of intrinsic activity and has been linked to both cognitive function and psychiatric disorders. Yet its connection to the brain’s neurochemical and cellular architecture remains poorly understood. By integrating molecular imaging with neuroimaging datasets, we show that the spatial distribution of cell types, neurotransmitter systems, and mitochondrial phenotypes is systematically related to brain entropy. We observed significant differences in entropy correlations: between healthy controls and individuals with schizophrenia for the mu-opioid and dopamine D1 receptors, and between controls and individuals with bipolar disorder for the norepinephrine transporter (NAT) and N-methyl-D-aspartate (NMDA) receptor. No significant differences emerged between schizophrenia and bipolar disorder in the neurotransmitter domain. At the cellular and metabolic level, both control-schizophrenia and control-bipolar comparisons revealed widespread alterations involving most mitochondrial markers, glial cells, and inhibitory neurons. These patterns point to disruptions in energy metabolism, neuroinflammatory processes, and inhibitory regulation within the clinical groups. Overall, the findings indicate that brain entropy is not randomly distributed but is closely tied to specific neurochemical systems and cellular features. This systems-level view helps explain how the complexity of brain activity arises from molecular architecture, and how it is altered in psychiatric disorders. It also provides a biological foundation for understanding brain entropy in the context of psychosis.

## 1 Introduction

Brain entropy, viewed as a feature of complex dynamical systems such as the primate brain, measures the rate at which information is generated over time. A widely used estimator of entropy-related complexity is sample entropy (SampEn). SampEn quantifies the unpredictability of a time series by assessing the probability of repeating patterns, given a prior probabilistic distribution [1]. It has been broadly applied in neuroimaging and physiological research to evaluate signal complexity [2, 3, 4].

In addition to entropy measures obtained from resting-state fMRI (rsfMRI), analyzing the spatial distribution of neurotransmitter systems and cell types is crucial for understanding brain function [5, 6, 7, 8]. Neurotransmitter maps, for instance, capture the density and organization of major neurochemical systems that support cognition and behavior. These maps not only characterize typical brain activity but also demonstrate how altered signaling contributes to psychiatric disorders such as schizophrenia (SZ) and bipolar disorder (BP) [9, 10].

Examining the distribution of glial, neuronal, and mitochondrial populations provides further insight into the brain’s cellular architecture [6, 7, 8]. Relating these cellular and neurochemical features to brain entropy helps clarify how disruptions in organization underlie neural abnormalities observed in psychiatric conditions, potentially driving entropy alterations themselves.

Combining neurotransmitter and cell-type maps with brain entropy analysis provides a powerful framework for investigating the biological mechanisms that underlie psychosis and other neuropsychiatric disorders. This multidimensional approach supports a more refined understanding of how changes in activity complexity relate to cellular and neurochemical disturbances. It also highlights potential biomarkers for diagnosis and points to new therapeutic targets.

## 2 Materials and Methods

### 2.1 Participants

A total of 1,432 participants were drawn from the Bipolar and Schizophrenia Network for Intermediate Phenotypes (BSNIP) consortium [11, 12]. The cohort comprised 640 healthy controls (NC), 470 individuals with SZ, and 322 with BP. Demographic and clinical characteristics of the participants are summarized in Table. 1.

**Table. 1.** Clinical and demographic information for the patient and healthy control groups.

|  | **SZ** | **BP** | **NC** |
| --- | --- | --- | --- |
| No. of Participants | 470 | 322 | 640 |
| Age (Mean ± SD) | 36.40 ± 12.13 | 34.30 ± 12.17 | 35.81 ± 12.28 |
| Sex (M/F) | 168/302 | 204/118 | 379/261 |
| PANSS Positive | 16.20 ± 5.77 | 12.27 ± 4.42 | N.A |
| PANSS Negative | 16.27 ± 6.39 | 11.71 ± 4.43 | N.A |
| PANSS General | 30.68 ± 9.34 | 27.88 ± 8.28 | N.A |
| PANSS Total | 63.13 ± 18.74 | 51.87 ± 14.88 | N.A |

\*PANSS stands for Positive and Negative Syndrome Scale.

An initial evaluation of the dataset revealed significant group differences in age and sex, based on two-sample t-tests applied to all pairwise comparisons. Detailed results of these tests are reported in Table. 2. To minimize potential confounding effects, age, sex, and other relevant covariates were regressed out in all subsequent analyses.

**Table. 2.**
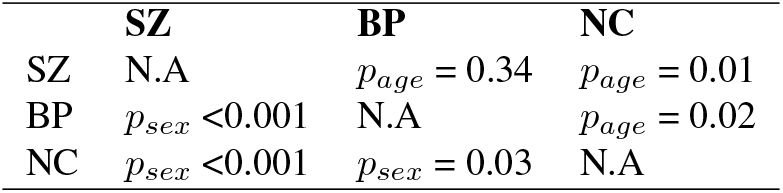
Statistical comparison results of age and sex across all pairwise group combinations. The upper triangle displays p-values for age comparisons, while the lower triangle shows p-values for sex.

|  | **SZ** | **BP** | **NC** |
| --- | --- | --- | --- |
| SZ | N.A | $p_{age} = 0.34$ | $p_{age} = 0.01$ |
| BP | $p_{sex} < 0.001$ | N.A | $p_{age} = 0.02$ |
| NC | $p_{sex} < 0.001$ | $p_{sex} = 0.03$ | N.A |

### 2.2 MRI Data Acquisition and Preprocessing

Our rsfMRI data underwent a rigorous preprocessing pipeline to ensure data quality and analytic reliability. The process began with strict quality control to retain only high-quality datasets. Each participant’s rsfMRI data were then processed using a standardized pipeline that included rigid-body motion correction, slice-timing correction, and distortion correction. The data were subsequently registered to a common MNI space, resampled to isotropic voxels of 3mm^3^, and smoothed with a Gaussian kernel having a full width at half maximum (FWHM) of 6mm.

### 2.3 Assessing Brain Signal Complexity via SampEn Measures

SampEn quantifies signal complexity by calculating the negative natural logarithm of the conditional probability that two sequences, similar for *m* consecutive points, remain similar at the next point within a tolerance *r* [1]. The tolerance *r* defines the maximum distance for two points to be considered similar and is commonly set to 0.2 × std(*X*), where std denotes the standard deviation of the time series *X*. Unlike related measures, SampEn excludes self-matches, producing more reliable results for shorter time series. In this study, we used *m* = 3 and *r* = 0.3, representing the embedding dimension and similarity tolerance, respectively. Notably, SampEn is always non-negative, as it is defined as the negative logarithm of a conditional probability.

For rsfMRI signals, a time series can be represented as a set of observations, *x*(*i*), where *i* denotes the time point and *x*(*i*) is the signal value at the corresponding voxel. Given a series of length *T*, a template vector of length *l* is defined as

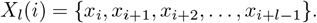

This vector captures a segment of the fMRI time series and serves to analyze the complexity of brain activity. Dissimilarity between two vectors, *X*_*l*_(*i*) and *X*_*l*_(*j*) with *i* ≠ *j*, is quantified using a distance function 𝒟 [*X*_*l*_(*i*), *X*_*l*_(*j*)], based on a selected threshold *r*. SampEn is then calculated as

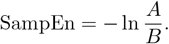

Here, *A* denotes the number of template vector pairs (*i, j*) for which the distance between their extended vectors of length *l* + 1, *X*_*l*+1_(*i*) and *X*_*l*+1_(*j*), is less than a tolerance *r*, i.e., 𝒟 [*X*_*l*+1_(*i*), *X*_*l*+1_(*j*)] *< r*. Similarly, *B* represents the number of pairs (*i, j*) where the distance between the original vectors of length *l, X*_*l*_(*i*) and *X*_*l*_(*j*), is below *r*, i.e., 𝒟 [*X*_*l*_(*i*), *X*_*l*_(*j*)] *< r*.

For fMRI time series, SampEn quantifies the regularity and complexity of brain activity over time. Lower SampEn values indicate more predictable activity, whereas higher values reflect greater complexity and less predictability. This metric is particularly useful for analyzing spontaneous brain activity during rsfMRI, as it captures the temporal structure and dynamic behavior of the brain signal.

### 2.4 Voxel-wise Correlation Between Brain Entropy and Positive and Negative Syndrome Scale (PANSS) Scores

Voxel-wise correlation analyses were conducted to investigate the relationship between regional brain entropy and symptom severity in individuals with SZ and BP. Clinical symptoms were measured using the PANSS, including subscale scores for positive, negative, and general psychopathology as independent predictors. Within each participant group, voxel-wise Pearson correlations were computed between entropy values and each PANSS subscale across all brain voxels contained in a predefined brain mask.

To account for multiple comparisons, voxel-wise p-values were corrected using the false discovery rate (FDR) method, with a significance threshold of *p <* 0.05. Voxels that survived FDR correction were considered statistically significant.

### 2.5 Spatial Correlation Analysis of Brain Entropy Alterations with Neurotransmitter Density Profiles and Cell-Type Distribution Maps

A set of 30 neurotransmitter and metabolic maps was selected, derived from positron emission tomography (PET), single-photon emission computed tomography (SPECT), and MRI, including arterial spin labeling-derived cerebral blood flow (ASL_CBF). These maps included receptor and transporter subtypes such as 5-hydroxytryptamine (serotonin, 5HT) receptors (5HT_1a_ [13, 14], 5HT_1b_ [13], 5HT_2a_ [13], 5HT_4_ [14], 5HT_6_ [15]), cannabinoid receptor type 1 (CB1) [16], cerebral blood flow (CBF) [17], dopamine receptors (D1 [18], D2 [19]), 6-[^18^F]fluoro-L-DOPA (FDOPA) [20], dopamine transporter (DAT) [21], noradrenaline transporter (NAT) [22], gamma-aminobutyric acid type A receptors (GABA_a_ [21], GABA_a5_ [23]), mu-opioid receptor (MU, MOR) [24], N-methyl-D-aspartate receptor (NMDA) [25], serotonin transporter (SERT) [13, 14, 26], metabotropic glutamate receptor 5 (mGluR5) [5], vesicular acetylcholine transporter (VAChT) [5], nicotinic acetylcholine receptor *α*4*β*2 (A4B2) [27], cerebral metabolic rate of glucose consumption (CMRglu) [28], cyclooxygenase-1 (COX1) [29], histone deacetylase (HDAC) [30], kappa-opioid receptor (KOR) [31], muscarinic acetylcholine receptor M1 (M1) [32], microglial marker (Microglia) [33], synaptic vesicle glycoprotein 2A (SV2A) [34], and vesicular monoamine transporter type 2 (VMAT2) [35]. These neurotransmitter maps are publicly available at https://github.com/netneurolab/hansen_receptors.

In addition to neurotransmitter and metabolic density maps, we incorporated 18 cell-type and 6 mitochondrial function distribution maps to further explore the biological basis of brain entropy alterations. These maps, derived from transcriptomic and histological data in healthy human brain tissue, capture the spatial distribution of glial, neuronal, and mitochondrial components. Glial maps [6] include astrocytes (Glia-Astro), endothelial cells (Glia-Endo), microglia (Glia-Micro), oligodendrocyte precursor cells (Glia-OPC), and mature oligodendrocytes (Glia-Oligo). Mitochondrial function [8] is represented by maps reflecting enzymatic activity and metabolic capacity, including Complex I (NADH), Complex II (succinate dehydrogenase, SDH), Complex IV (cytochrome c oxidase, COX), overall respiratory capacity, tissue density, and tissue-specific respiratory capacity. These maps are available at https://neurovault.org/collections/16418/. Neuronal cell-type maps [7] cover both excitatory and inhibitory sub-populations. Excitatory subtypes (Ex1–Ex7) are defined by their projection targets and laminar locations. Inhibitory interneurons are categorized based on molecular markers such as vasoactive intestinal peptide (VIP), somatostatin (SST), reelin (RELN), neuronal-derived neurotrophic factor (NDNF), cholecystokinin (CCK), parvalbumin (PVALB), and nitric oxide synthase (NOS). These cell-type and mitochondrial maps were selected to capture detailed cellular and metabolic het-erogeneity across the human cortex, providing key insights into potential mechanisms underlying altered entropy patterns observed in neuropsychiatric disorders.

### 2.6 Statistical Analysis

To quantify the magnitude of group differences in brain entropy between diagnostic groups, we computed voxel-wise Cohen’s *d* effect size maps. This analysis was performed across all voxels to produce spatial maps that reflect the extent of entropy alterations between groups. Positive *d* values indicate higher entropy in one group relative to the other, while negative values indicate the opposite. These spatial effect size maps offer a detailed view of how intrinsic brain activity complexity varies across psychosis-spectrum conditions and highlight brain regions most affected in each comparison.

Furthermore, Spearman correlation analysis was performed to investigate associations between brain entropy alterations, extracted using the brain atlas, and spatial distributions of neurotransmitter and cell-type values mapped to the same atlas. Comparisons were conducted across diagnostic groups. Spearman correlation coefficients, Fisher’s z-transformed, were calculated between z-scored brain entropy alteration maps and each selected neurotransmitter or cell-type distribution map.

## 3 Experimental Results

### 3.1 Brain Entropy, Neurotransmitter, and Cellular Expression Values in the Brain

As shown in Fig. 1, voxel-wise SampEn was first estimated from preprocessed resting-state fMRI data collected from a large cohort of subjects. These voxel-level estimates were then aggregated into regional values using a brain atlas (e.g., Neuromorphometrics). In parallel, 30 neurotransmitter density maps and 24 cellular distribution maps were processed. The cellular maps included various neuronal and glial subtypes, as well as mitochondrial markers. All maps were registered and resampled to MNI space to ensure alignment with the fMRI data. Regional values for neurotransmitter and cellular features were extracted using the same brain atlas, enabling consistent spatial comparisons across modalities.

**Fig. 1.**
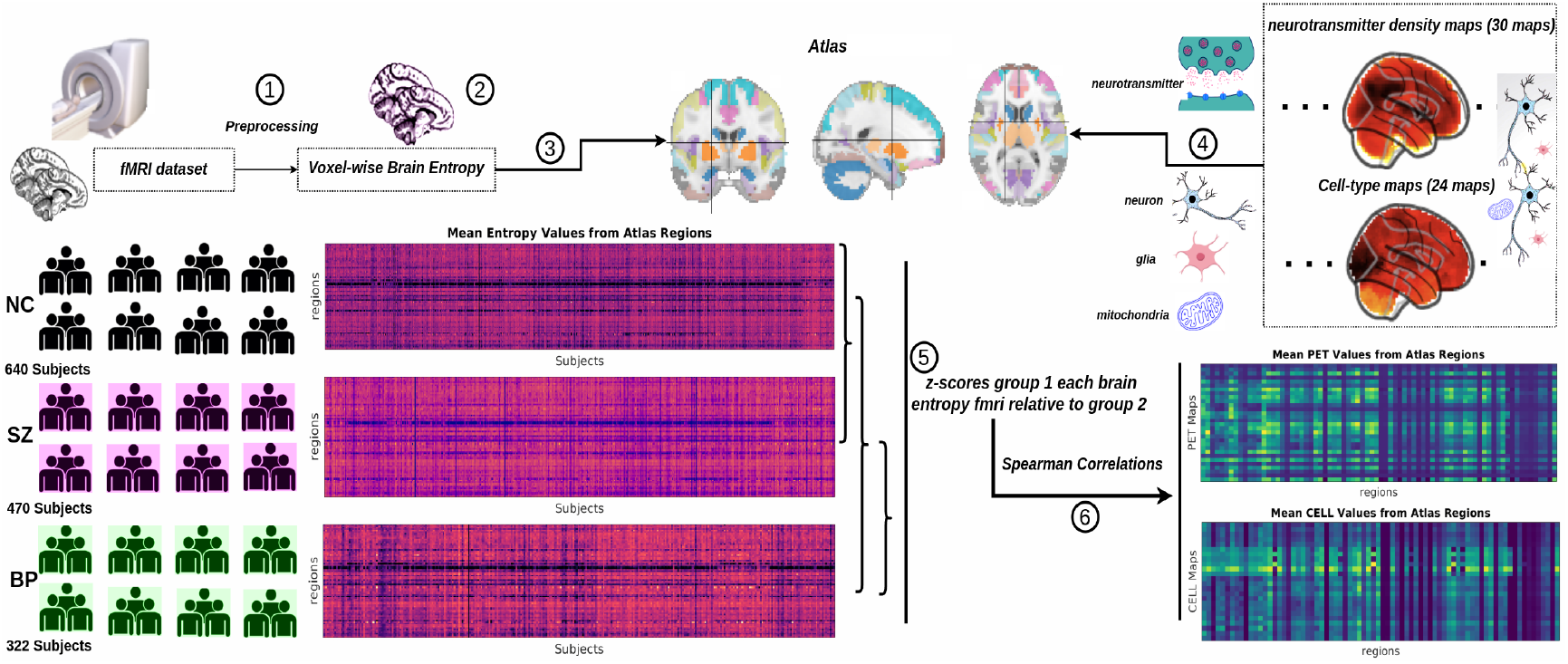
Analysis pipeline. Raw resting-state fMRI data from NC, SZ, and BP participants were preprocessed and used to compute voxel-wise brain entropy using SampEn. Mean brain entropy values were then extracted for each group using a brain atlas. In parallel, mean values were obtained from 30 neurotransmitter maps (derived from PET) and 24 cell-type expression maps, which include 18 neuronal and glial subtypes and 6 mitochondrial function maps. Group differences in brain entropy were quantified by computing z-scores (e.g., NC vs. SZ, NC vs. BP, SZ vs. BP). Spearman correlation analyses were subsequently performed between group-level brain entropy alterations and molecular (PET) and cellular (cell-type) maps. Correlations were FDR-corrected for multiple comparisons at *p <* 0.05. This pipeline highlights the integration of functional, molecular, and cellular data to identify biologically relevant group differences in brain entropy.

### 3.2 Voxel-wise Brain Entropy Effect Size Maps Across Psychosis

We quantified differences in brain entropy across diagnostic groups by computing voxel-wise effect size maps using Cohen’s *d*. Specifically, pairwise comparisons were performed between NC and SZ, NC and BP, and SZ and BP. As illustrated in Fig. 2, we observed that the orbitofrontal cortex exhibits higher SampEn in SZ, which reflects increased complexity and irregularity in neural activity. This heightened entropy indicates less predictability in the temporal patterns of brain activity, suggesting that the neural signals in this region become more disordered or variable. These irregularities in neural activity may be linked to impairments in cognitive functioning. Notably, this finding closely aligns with prior clinical studies [36], which have documented comparable alterations in the orbitofrontal cortex among individuals with SZ, thereby strengthening the evidence for its critical role in the pathophysiology of SZ and related psychiatric conditions, including BP. In the case of BP, we also observed pronounced increases in sample entropy within the visual cortex, suggesting heightened neural complexity and dysregulation in sensory processing regions that may reflect disorder-specific neural dynamics.

**Fig. 2.**
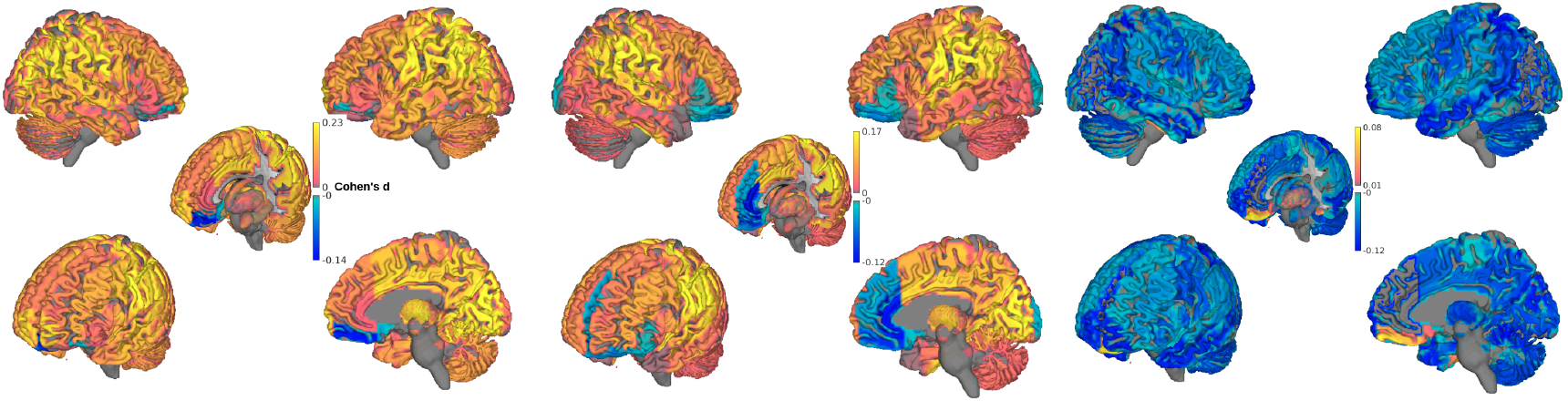
Effect size maps across psychosis. Cohen’s *d* effect sizes of brain entropy were computed for each pairwise group comparison: NC vs. SZ, NC vs. BP, and SZ vs. BP. The resulting effect sizes were projected onto the brain surface to visualize regional patterns of group differences. Positive values indicate higher entropy in the first group of the comparison, while negative values indicate higher entropy in the second group. All maps show only regions surviving FDR correction at *p <* 0.05. Color scales represent the magnitude and direction of Cohen’s *d*, highlighting regions with the strongest effects across groups.

In addition, given the substantial clinical overlap between SZ and BP, distinguishing between these two conditions remains a significant challenge. However, our SampEn analysis revealed that the orbitofrontal cortex exhibits markedly stronger differences between SZ and BP, suggesting that this region may serve as a promising neural marker for differentiating these clinically similar disorders.

### 3.3 Voxel-wise Associations Between Brain Entropy and PANSS Subscales

Voxel-wise correlation analyses revealed distinct spatial patterns of association between brain entropy and PANSS symptom dimensions in the SZ and BP groups, as illustrated in Fig. 3.

**Fig. 3.**
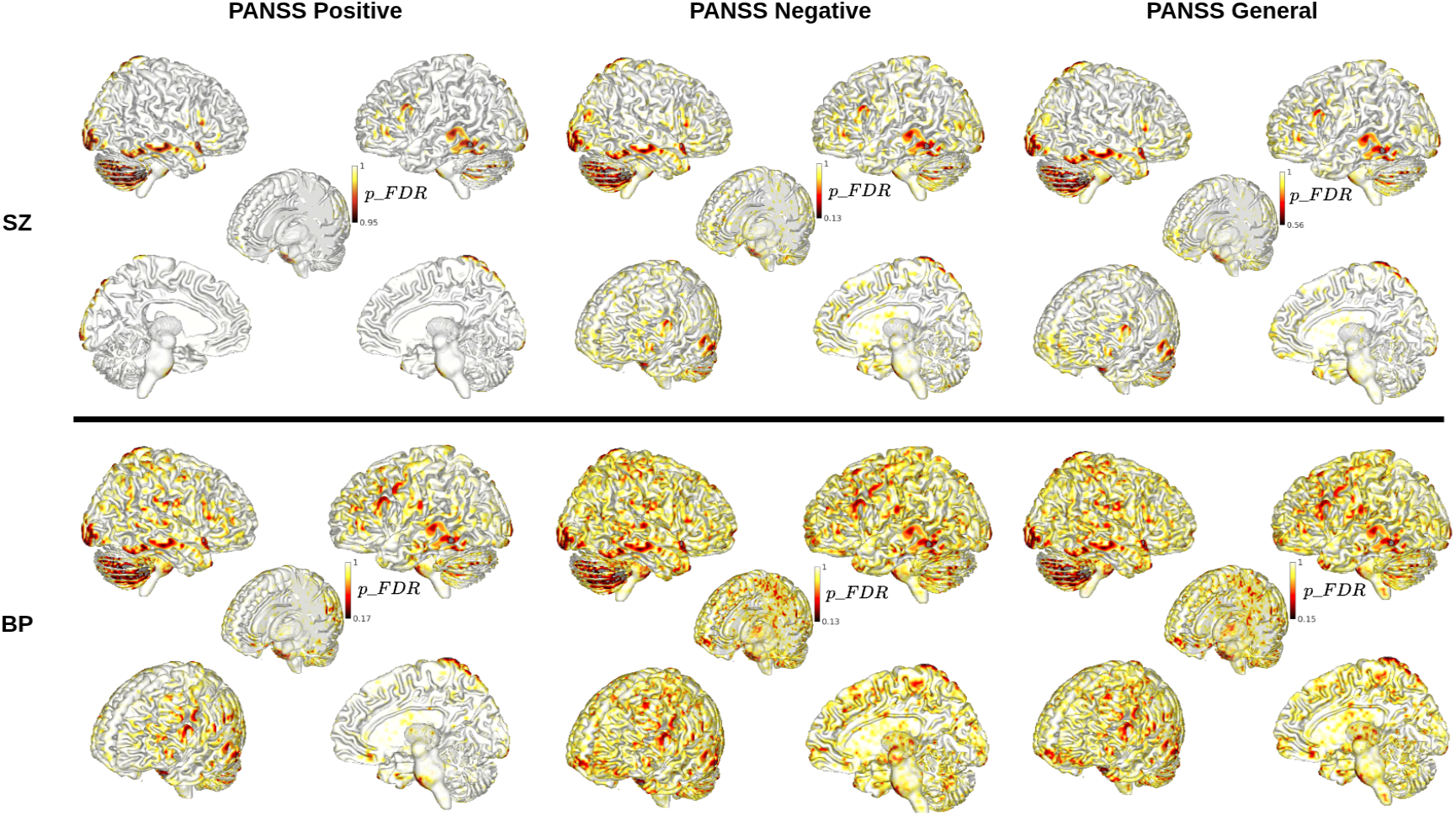
Brain regions where entropy correlates with PANSS scores. Voxel-wise correlations between brain entropy and PANSS subscale scores (positive, negative, and general psychopathology) were computed separately for the SZ and BP cohorts. Only voxels showing significant associations after FDR correction (*p <* 0.05, FDR-corrected) are displayed. Color scales represent the strength and direction of the correlation, with positive values indicating higher entropy associated with higher symptom severity and negative values indicating the opposite. This map highlights brain regions where local entropy reliably reflects clinical symptom severity.

In the SZ group, significant correlations were observed between brain entropy and PANSS positive symptom scores, primarily in the occipital visual cortex, temporal lobes, and cerebellum. Additional clusters showing significant associ-ations with PANSS negative and general psychopathology scores were located in the parietal and frontal cortices.

In contrast, the BP group showed both similarities and differences in the spatial pattern of correlations between brain entropy and PANSS scores. Similar to the SZ group, significant associations were observed in the occipital visual cortex, temporal lobes, and cerebellum. However, the BP group exhibited a more widespread distribution of significant correlations across the whole brain. The most prominent difference was the location of peak correlations, which were concentrated in the prefrontal cortex, distinguishing the BP profile from the more occipito-temporal and cerebellar involvement observed in SZ.

### 3.4 Brain Entropy Alterations in Relation to Neurotransmitter and Metabolic Distributions

As shown in Fig. 4***A***, the 30 neurotransmitter maps were plotted to visualize the spatial distribution of neurotransmitter expression across the brain. In Fig. 4***B***, we show SampEn alterations showed significant spatial correlations with several neurotransmitter distributions across diagnostic comparisons. In patients with SZ relative to NC, SampEn alterations were significantly associated with the spatial distribution of dopamine receptor (D1) (*r* = −0.044, *p* = 0.030), mu-opioid receptors (MOR; *r* = −0.052, *p* = 0.015), and MU receptors (*r* = −0.051, *p* = 0.014), all surviving FDR correction at *p <* 0.05. This result is particularly interesting and aligns well with previous studies, which have demonstrated that in SZ, both dopamine and mu-opioid receptor systems play crucial roles, with consistent evidence indicating alterations in their function and availability [37, 38].

**Fig. 4.**
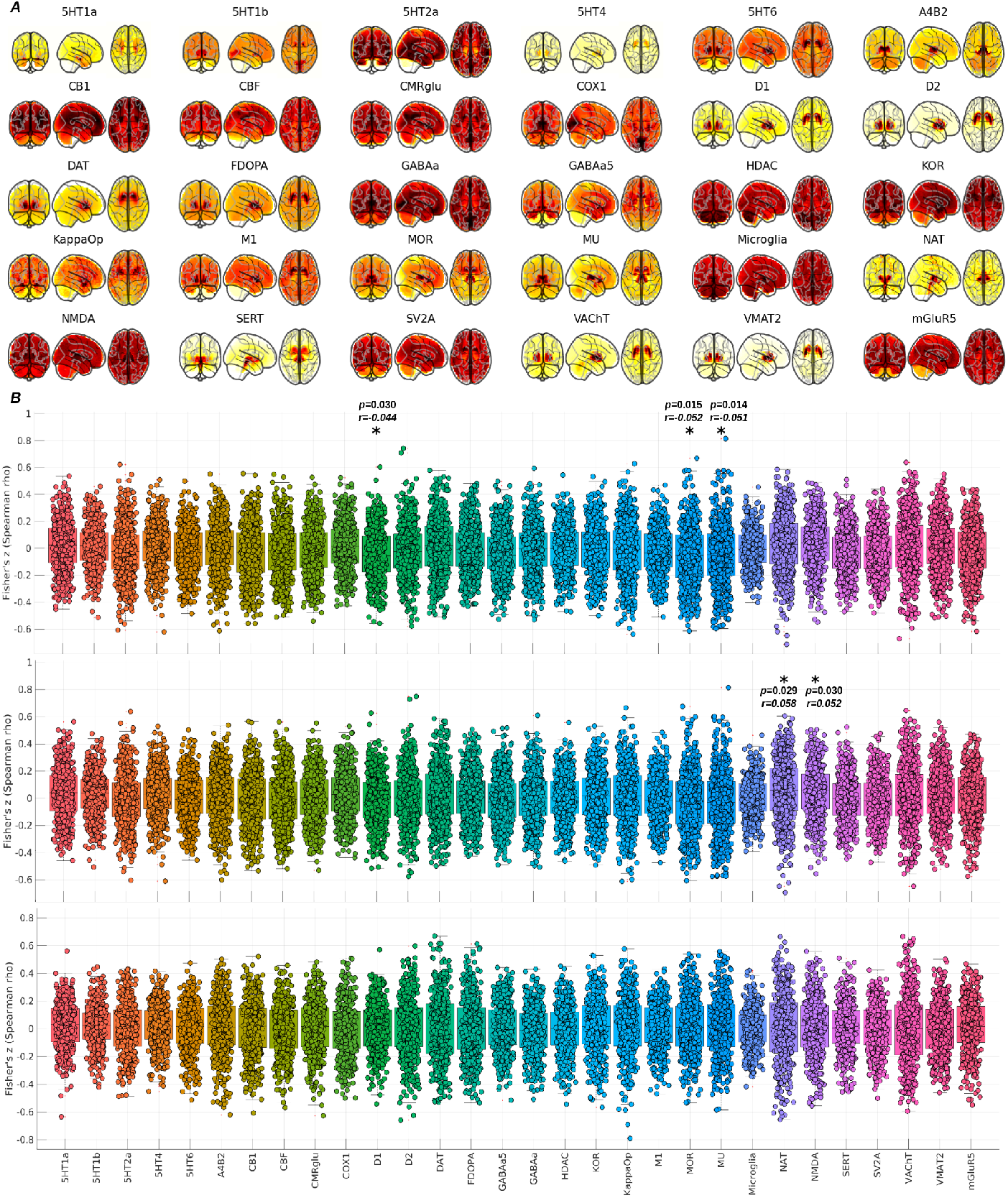
Spatial correlation analyses between brain entropy alterations and neurotransmitter and metabolic distributions. ***A***. Thirty neurotransmitter and metabolic density maps used in the analysis. ***B***. Spatial correlation results based on the Neuromorphometrics atlas. From top to bottom, rows represent correlations between brain entropy alterations and neurotransmitter or metabolic maps for NC vs. SZ, NC vs. BP, and SZ vs. BP. Significant associations after FDR correction (*p <* 0.05) are indicated by *. This panel highlights some molecular systems that are most strongly associated with group-level entropy alterations.

In BP patients compared with NC, SampEn alterations were significantly associated with the spatial distribution of noradrenaline transporters (NAT; *r* = 0.058, p = 0.029) and NMDA receptors (*r* = 0.052, p = 0.030), with all results corrected-FDR at p < 0.05. This pattern supports prior evidence linking these receptors to the pathophysiology of BP [39, 40]. Altered receptor function may contribute to clinical symptoms and disease progression, highlighting the neurobiological basis of BP.

However, no significant correlations between SampEn alterations and neurotransmitter distributions were observed in the SZ versus BP comparison.

### 3.5 Brain Entropy Alterations in Relation to Cellular-Type Maps

As shown in Fig 5***A***, the spatial distribution of cell-type expression across the brain was visualized using 24 cell-type distribution maps, encompassing neuronal, glial, and mitochon-drial cell types. In Fig. 5***B***, alterations in SampEn demonstrated significant spatial correlations with several cell-type distributions across diagnostic groups. In patients with SZ compared to NC, SampEn alterations were significantly associated with the spatial distribution of mitochondria and glial cells, particu-larly astrocytes (Glia-Astro; *r* = −0.030, *p* = 0.024), as well as inhibitory neurons, including neuron-In1-VIP-RELN-NDNF-L12 (*r* = −0.038, *p* = 0.019) and neuron-In4-RELN-NDNF-L13 (*r* = 0.039, *p* = 0.019). These findings suggest that the altered brain signal complexity in SZ may be linked to disruptions in energy metabolism, neuroinflammatory processes, and inhibitory neurotransmission. Meanwhile, these findings provide a biological foundation for brain entropy, supporting its potential utility as a proxy marker for underlying cellular dysfunction and as a quantitative index for characterizing neuropathological changes in SZ.

**Fig. 5.**
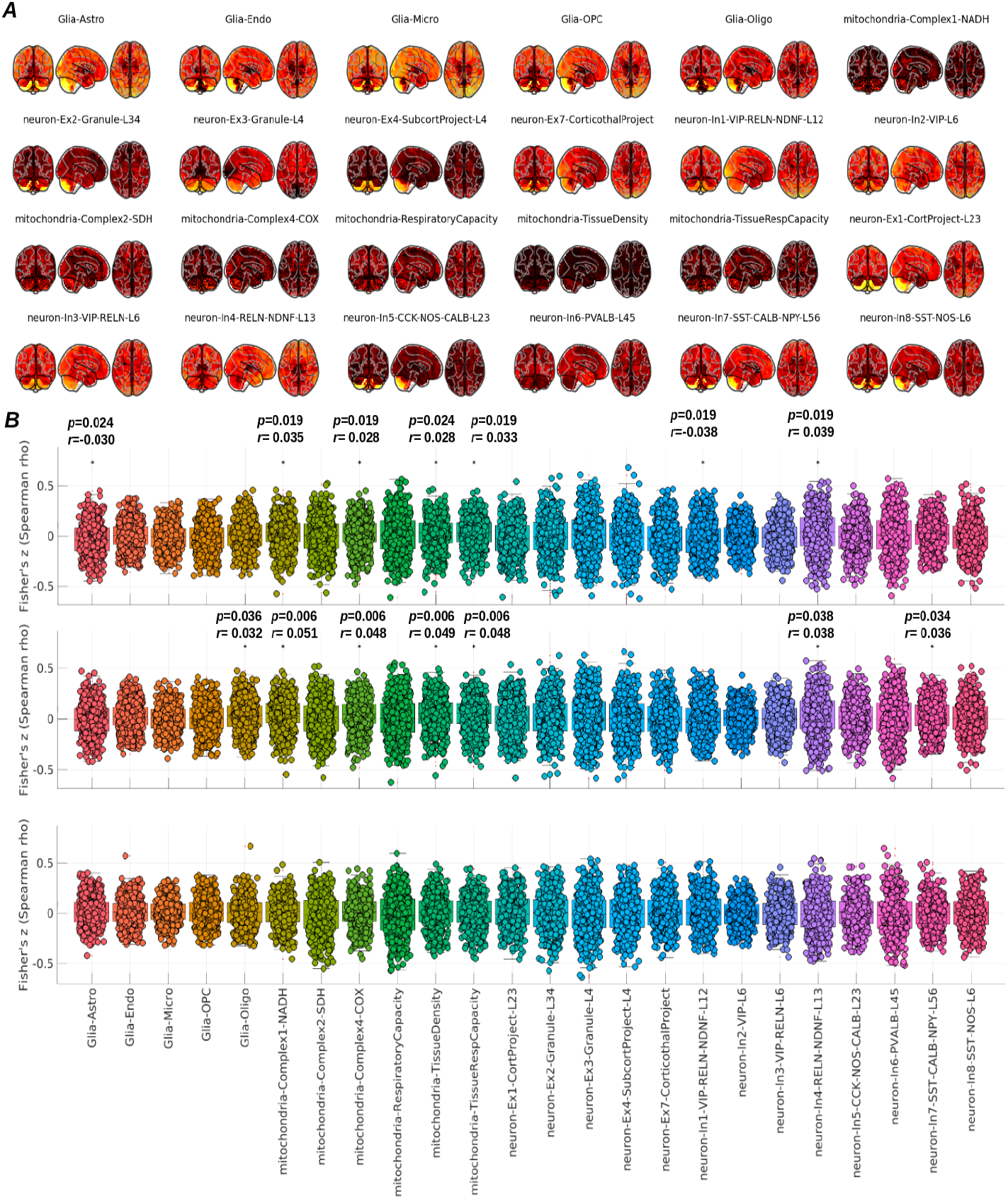
Results of spatial correlation analyses between brain entropy alterations and cell-type distribution. ***A***. Twenty-four cell-type spatial maps. ***B***. Spatial correlation results based on the Neuromorphometrics atlas. From top to bottom, rows display correlations between brain entropy alterations and cell-type maps for NC vs. SZ, NC vs. BP, and SZ vs. BP. Each brain entropy alteration map was correlated with the cell-type maps indicated on the x-axis, with different colors representing each cell map. Significant associations are marked by an asterisk (*; *p* < 0.05, FDR-corrected).

In addition to findings in patients with SZ, patients with BP relative to NC also exhibited a similar pattern, in which SampEn alterations were significantly correlated with the spatial distribution of most mitochondrial cell types and neuron-In4-RELN-NDNF-L13 (*r* = 0.038, *p* = 0.038). This common pattern suggests that both SZ and BP may involve disruptions in energy metabolism at the cellular level. Furthermore, SampEn alterations in BP were also significantly associated with the spatial distribution of oligodendrocytes (Glia-Oligo; *r* = 0.032, *p* = 0.036), indicating potential involvement of myelination-related processes. These results further imply that SZ and BP share overlapping pathophysiological features, particularly related to mitochondrial dysfunction, neuroinflammatory activity, and inhibitory neuronal regulation.

Furthermore, no significant correlations between SampEn alterations and cell-type distributions were observed in the SZ versus BP comparison.

### 3.6 Identifying Key Neurotransmitter and Cell Type Associations with SampEn Alterations

As shown in Fig. 6, to facilitate a better comparison of the relationships between SampEn alterations and the spatial distributions of neurotransmitters and cell types, we plotted all FDR-corrected *p*-values in ascending order. Features exhibiting significant associations (*p* < 0.05) were highlighted and labeled accordingly. This visualization enhances the comparison of effect strengths across different features, enabling the identification of the most strongly associated neurotransmitters and cell types contributing to entropy alterations.

**Fig. 6.**
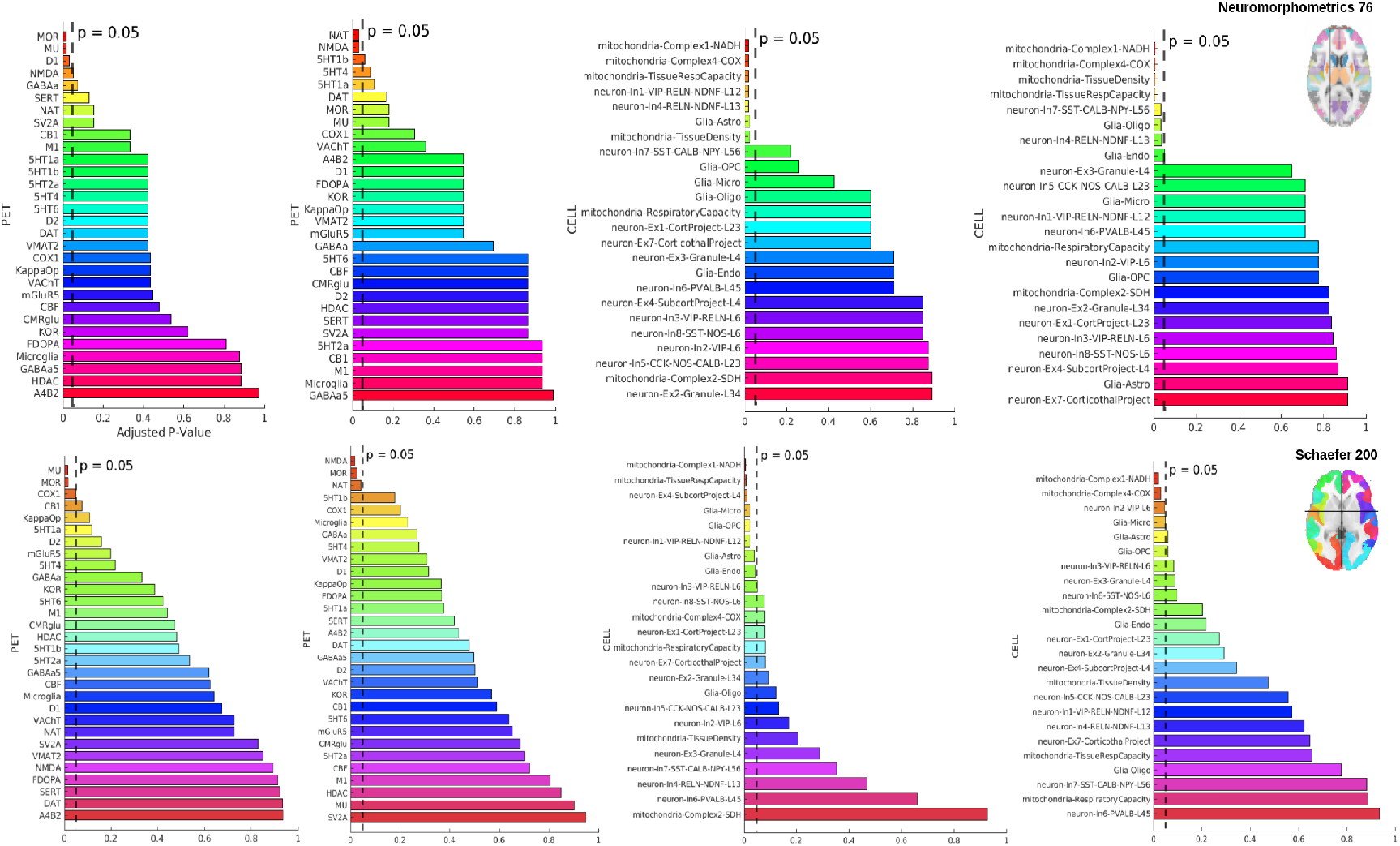
Associations of SampEn alterations with neurotransmitter and cell-type expression. From left to right, the figure shows spatial correlations between SampEn alterations and the distributions of neurotransmitters and cell types: NC vs. SZ in the first and third columns, and NC vs. BP in the second and fourth columns. The top row presents results based on the Neuromorphometrics atlas, which defines anatomically based, low-resolution brain regions. The bottom row shows corresponding results using the Schaefer 200-parcel atlas, a higher-resolution functional parcellation. Significant associations after FDR correction (*p* < 0.05) are indicated by dotted lines. Color scales represent p-values, indicating the strength of the associations.

To further validate our results, we employed a higher-resolution brain atlas (e.g., Schaefer 200 [41]) and applied the same analysis pipeline. The corresponding results are presented in Fig. 6. The findings were largely consistent with those obtained using the original atlas, offering additional evidence for the robustness of our initial analysis. This supplementary validation further strengthens the reliability and generalizability of our conclusions.

## 4 Limitations of the Study

One limitation of this study is its reliance on predefined anatomical or functional brain atlases. Incorporating totally data-driven or hybrid atlases, such as NeuroMark [42], could further enhance the precision and interpretability of future analyses. Second, the analysis does not incorporate transcriptomic data, which could provide a more comprehensive understanding of gene expression patterns and their relationship to brain entropy alterations. This limitation is primarily due to the lack of comprehensive voxel-wise transcriptomic maps covering the entire brain, which constrains the feasibility of such analyses within our current framework. Future research could address this by investigating these relationships at the functional connectivity level, where transcriptomic data may be more readily integrated. Additionally, examining functional connectivity differences between individuals with schizophrenia and bipolar disorder at this level would also be valuable for further elucidating disorder-specific neurobiological mechanisms. Third, the study focuses primarily on pairwise interactions between SampEn alterations and cell-type distributions, and does not explore higher-order interactions [43], which could reveal more complex relationships and offer deeper insights into the brain’s cellular architecture. Fourth, the analysis of neurotransmitter and cell-type distributions is constrained by the current resolution and specificity of available maps, which may not fully capture the diversity and dynamic changes of these populations across different brain regions. Finally, this study only examines static brain entropy, whereas dynamic brain entropy might offer more nuanced insights into the temporal fluctuations and network interactions that underlie brain activity.

In summary, these limitations highlight the need for future studies to incorporate data-driven brain atlases, transcriptomic data, higher-order network interactions, and dynamic entropy measures to gain a deeper understanding of the neurobiological mechanisms underlying disorders such as schizophrenia and bipolar disorder.

## 5 Conclusion

This study bridges the gap between brain entropy and the molecular and cellular architecture of the brain, providing a deeper understanding of psychosis. Moving beyond traditional macro-level approaches, we now explore the intricate relation-ships at the neurotransmitter and cellular levels. This shift offers a more nuanced perspective on the biological mechanisms underlying psychosis, enhancing our understanding of its manifestation at both systemic and cellular scales. Significant entropy correlations were found in psychosis, linked to several neurotransmitter systems. Additionally, we observed disruptions in energy metabolism, neuroinflammation, and inhibitory neuronal regulation, particularly at the cellular level, in mitochondrial markers. These findings highlight that brain entropy is systematically connected to specific neurochemical systems and cellular features, offering new insights into the biological mechanisms of psychosis. Furthermore, this study gives entropy a biological underpinning, demonstrating its relevance beyond its traditional roots in physics.

## 6 Ethics Statement

This study utilized data from the Bipolar-Schizophrenia Network on Intermediate Phenotypes (BSNIP) dataset [11, 12]. All data used were previously collected and fully de-identified in accordance with the ethical standards set by the original data-collecting institutions. The BSNIP study received approval from the institutional review boards (IRBs) of all participating sites, and written informed consent was obtained from all participants at the time of data collection. The current analysis was conducted using only de-identified data and did not involve any direct interaction with human participants.

## 7 Data and Code Availability Statement

The fMRI dataset is not publicly available due to licensing restrictions but can be requested from the NIH Data Archive at: https://nda.nih.gov/edit_collection.html?id=2274. Neurotransmitter maps used in this study are accessible at: https://github.com/netneurolab/hansen_receptors, while mitochondrial maps can be obtained via NeuroVault: https://neurovault.org/collections/16418/. Receptor analysis was performed using the JuSpace toolbox, available at: https://github.com/juryxy/JuSpace, and voxel-wise brain entropy was computed with the Brain Entropy Toolbox (BENtbx), accessible at: https://github.com/zewangnew/BENtbx.

## 9 Authorship Contribution Statement

Qiang Li: Writing – original draft, Visualization, Validation, Software, Methodology, Investigation, Formal analysis. Jingyu Liu: Writing – review & editing, Funding acquisition, Resources, Methodology. Vince D. Calhoun: Writing – review & editing, Funding acquisition, Resources, Methodology.

## 10 Acknowledgements

This work was partially supported by NSF grant 2112455, and the NIH grant R01MH123610.

